# Epigenetic prediction of complex traits and death

**DOI:** 10.1101/294116

**Authors:** Daniel L McCartney, Anna J Stevenson, Stuart J Ritchie, Rosie M Walker, Qian Zhang, Stewart W Morris, Archie Campbell, Alison D Murray, Heather C Whalley, Catharine R Gale, David J Porteous, Chris S Haley, Allan F McRae, Naomi R Wray, Peter M Visscher, Andrew M McIntosh, Kathryn L Evans, Ian J Deary, Riccardo E Marioni

**Affiliations:** Centre for Genomic and Experimental Medicine, Institute of Genetics and Molecular Medicine, University of Edinburgh, Edinburgh, EH4 2XU; Centre for Cognitive Ageing and Cognitive Epidemiology, University of Edinburgh, Edinburgh, EH8 9JZ; Department of Psychology, University of Edinburgh, Edinburgh, EH8 9JZ; Institute for Molecular Bioscience, University of Queensland, Brisbane, QLD, Australia; Aberdeen Biomedical Imaging Centre, Lilian Sutton Building, University of Aberdeen, Foresterhill, Aberdeen AB25□2ZD, UK; Division of Psychiatry, University of Edinburgh, Royal Edinburgh Hospital, Edinburgh, EH10 5HF, UK; MRC Human Genetics Unit, Institute of Genetics and Molecular Medicine, University of Edinburgh, Edinburgh, EH4 2XU

**Keywords:** DNA methylation, polygenic scores, prediction, ageing, mortality

## Abstract

**Background:** Genome-wide DNA methylation (DNAm) profiling has allowed for the development of molecular predictors for a multitude of traits and diseases. Such predictors may be more accurate than the self-reported phenotypes, and could have clinical applications. Here, penalised regression models were used to develop DNAm predictors for body mass index (BMI), smoking status, alcohol consumption, and educational attainment in a cohort of 5,100 individuals. Using an independent test cohort comprising 906 individuals, the proportion of phenotypic variance explained in each trait was examined for DNAm-based and genetic predictors. Receiver operator characteristic curves were generated to investigate the predictive performance of DNAm-based predictors, using dichotomised phenotypes. The relationship between DNAm scores and all-cause mortality (n = 214 events) was assessed via Cox proportional-hazards models.

**Results:** The DNAm-based predictors explained different proportions of the phenotypic variance for BMI (12%), smoking (60%), alcohol consumption (12%) and education (3%). The combined genetic and DNAm predictors explained 20% of the variance in BMI, 61% in smoking, 13% in alcohol consumption, and 6% in education. DNAm predictors for smoking, alcohol, and education but not BMI predicted mortality in univariate models. The predictors showed moderate discrimination of obesity (AUC=0.67) and alcohol consumption (AUC=0.75), and excellent discrimination of current smoking status (AUC=0.98). There was poorer discrimination of college-educated individuals (AUC=0.59).

**Conclusions:** DNAm predictors correlate with lifestyle factors that are associated with health and mortality. They may supplement DNAm-based predictors of age to identify the lifestyle profiles of individuals and predict disease risk.

**List of abbreviations:**

DNAm: DNA methylation
BMI: Body mass index
AUC: Area under the curve
CpG: Cytosine phosphate Guanine dinucleotide
EWAS: Epigenome-wide association study
GS:SFHS: Generation Scotland: The Scottish family health study
LBC1936: Lothian birth cohort 1936
LASSO: Least absolute shrinkage and selector operator
HR: Hazard ratio
CI: Confidence interval
STRADL: Stratifying resilience and depression longitudinally

## Background

DNA-based predictors of health and lifestyle have potential uses in both clinical and non-clinical contexts. For example, biological predictors of smoking status and alcohol consumption may provide more accurate measurements than self-report, thereby improving disease prediction and risk stratification [1]. Here, using whole blood-derived samples, we develop novel DNA methylation-based predictors of alcohol consumption, smoking status, body mass index (BMI), and educational attainment and relate them to both a health outcome (mortality) and lifestyle characteristics in an independent cohort.

DNA methylation (DNAm) is a commonly-studied epigenetic modification characterised by chemical changes to DNA – typically at a Cytosine-phosphate-Guanine (CpG) nucleotide base pairing [2]. These modifications are dynamic, tissue- and cell-specific [3], are involved in gene regulation, and can be influenced by both genes and the environment [4].

Through large meta-analysis projects, methylation signals at individual CpG sites have been associated with educational attainment, smoking, alcohol consumption, and BMI [5, 6, 7, 8, 9]. Such studies have also used methylation predictors (from a combination of CpG sites) to predict the phenotype of interest in independent cohorts. For example, 7% of the variance in BMI and 2% of the variance in educational attainment can be explained by their respective predictors [5, 10]. Studies have also combined genetic risk scores into their prediction models, showing that the DNAm predictors contribute independently to the variance explained in BMI and C-reactive protein levels [9, 11]. Moreover, single CpG sites and DNAm predictors have been linked to lung cancer/mortality [12], and cardiometabolic traits [7, 11].

There are, however, several limitations to existing studies. First, the CpG weights for the predictors are derived separately for each CpG, which does not account for their inter-correlations. Second, large samples are required to generate precise weights. This has meant conducting meta-analyses with data from heterogeneous populations where different quality control metrics have been applied. Third, the CpG prediction weights are typically based on Z-scores rather than effect sizes, that is, the trait was modelled as the predictor with the CpG as the outcome in the epigenome-wide association studies (EWASs). These Z-score weights are equivalent to modelling by p-values, which don’t account for the magnitude of the CpG-trait association. Fourth, arbitrary significance threshold cut-offs are used to select the number of CpGs used in each predictor rather than training a predictor on an optimised set of CpGs.

Here, we model all CpGs simultaneously in a single large cohort of over 5,000 individuals. We model the traits of interest as the outcomes and the CpGs as the predictors and train optimised predictors using penalised regression methods. We then apply these predictors to an independent cohort study of over 900 individuals to determine 1: the proportion of variance the DNAm predictors explain in the outcomes; 2: the extent to which these proportions are independent from the contribution of genetics; 3: the accuracy with which the DNAm predictors can identify obese individuals, college-educated individuals, heavy drinkers, and current smokers if provided with a random DNA sample from the population; and 4: the extent to which they can predict health outcomes, such as mortality and if they do so independently from the phenotypic measure.

## Results

Summary information on the four phenotypes in both the training (GS) and test (LBC1936) datasets is presented in **Table 1**. LBC1936 is an older cohort than GS (mean age 70 vs 48 years), with a more even gender balance (51% vs 39% male). LBC1936, when compared with GS participants, had around 2 fewer years of education, were of similar mean BMI (both cohort means were ~27kg/m^2^), drank slightly less alcohol (median difference of 3 units per week), and had a lower ratio of current to never smokers (20% vs 27%).

**Table 1:**
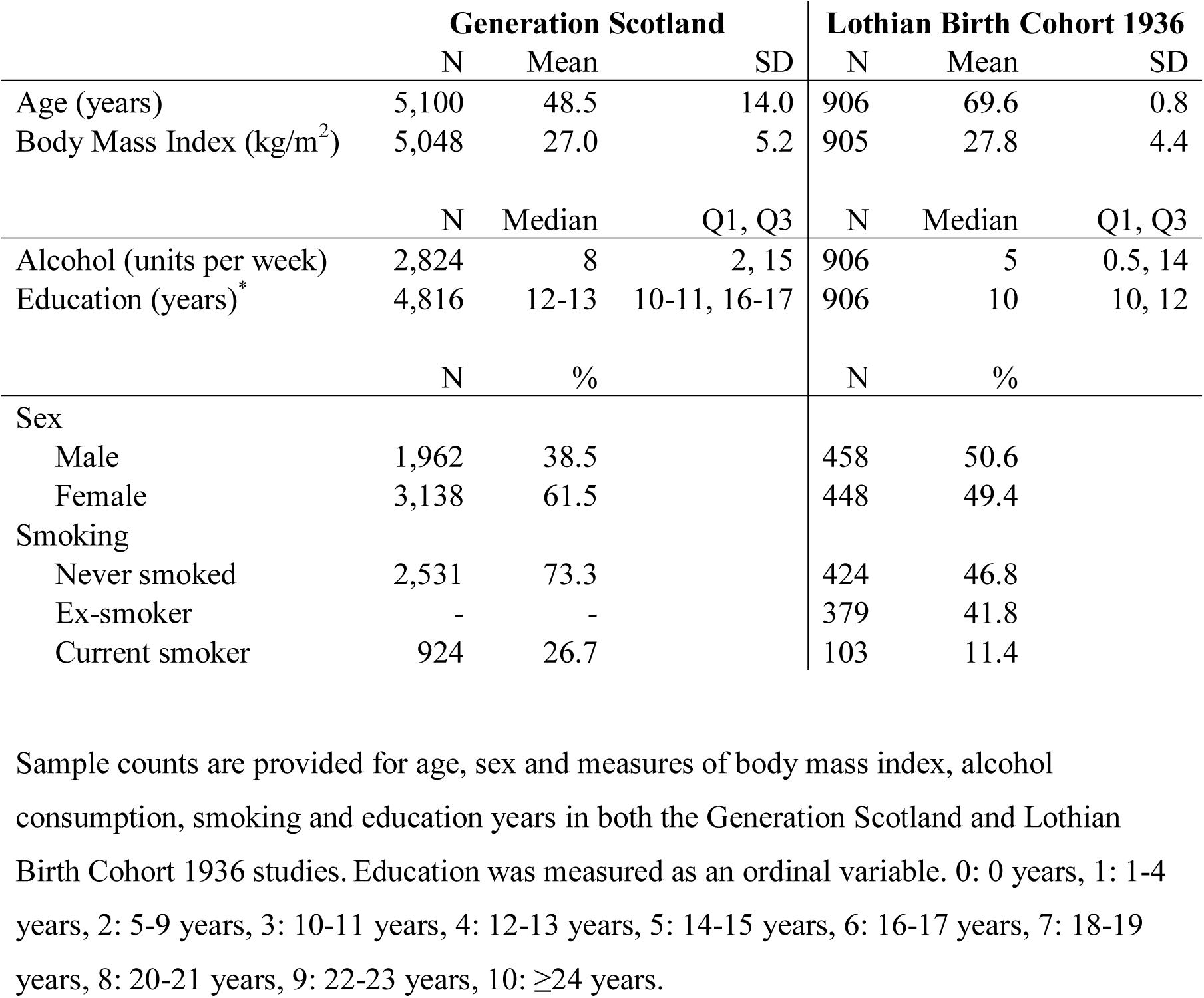
Summary of the Generation Scotland and Lothian Birth Cohort 1936 studies

|  | Generation Scotland |  |  | Lothian Birth Cohort 1936 |  |  |
| --- | --- | --- | --- | --- | --- | --- |
|  | N | Mean | SD | N | Mean | SD |
| Age (years) | 5,100 | 48.5 | 14.0 | 906 | 69.6 | 0.8 |
| Body Mass Index (kg/m<sup>2</sup>) | 5,048 | 27.0 | 5.2 | 905 | 27.8 | 4.4 |
|  | N | Median | Q1, Q3 | N | Median | Q1, Q3 |
| Alcohol (units per week) | 2,824 | 8 | 2, 15 | 906 | 5 | 0.5, 14 |
| Education (years)* | 4,816 | 12-13 | 10-11, 16-17 | 906 | 10 | 10, 12 |
|  | N | % |  | N | % |  |
| Sex |  |  |  |  |  |  |
| Male | 1,962 | 38.5 |  | 458 | 50.6 |  |
| Female | 3,138 | 61.5 |  | 448 | 49.4 |  |
| Smoking |  |  |  |  |  |  |
| Never smoked | 2,531 | 73.3 |  | 424 | 46.8 |  |
| Ex-smoker | - | - |  | 379 | 41.8 |  |
| Current smoker | 924 | 26.7 |  | 103 | 11.4 |  |

Sample counts are provided for age, sex and measures of body mass index, alcohol consumption, smoking and education years in both the Generation Scotland and Lothian Birth Cohort 1936 studies. Education was measured as an ordinal variable. 0: 0 years, 1: 1-4 years, 2: 5-9 years, 3: 10-11 years, 4: 12-13 years, 5: 14-15 years, 6: 16-17 years, 7: 18-19 years, 8: 20-21 years, 9: 22-23 years, 10: $\geq 24$ years.

The LASSO regressions returned predictors based on 1,099 (BMI), 287 (smoking), 371 (alcohol), and 281 CpGs (educational attainment). The regression weights for the predictors are shown in **Supplementary Tables 1-4**. DNAm predictors for the four variables were created in LBC1936 at the baseline wave, at a mean age of approximately 70 years (n=906).

Correlations between the phenotypic measures in GS:SHFS are presented in **Supplementary Table 5**. Correlations between the phenotypic measures and DNAm predictors in LBC1936 are presented in **Supplementary Table 6**. Small correlations (r < 0.2) were seen between the phenotypes, and also between the DNAm predictors. An exception was the DNAm smoking:DNAm education correlation (r = −0.54); the phenotypic smoking:DNAm education association was of a similar magnitude (r = −0.44).

### DNAm predictors explain phenotypic variation

Age and sex-adjusted linear regression models showed that the DNAm predictors, which were developed in GS, explained 12.2% of the variance in BMI, 12.5% in alcohol consumption, 60.6% in smoking, and 2.6% in education in LBC1936 (**Table 2** and **Figure 1**). The corresponding polygenic scores explained 10.3% of the variance in BMI, 0.7% in alcohol consumption, 2.8% in smoking, and 4% in education. Models including both the DNAm predictor and the polygenic score explained the most variance in each trait: 19.5% in BMI, 12.9% in alcohol consumption, 61.0% in smoking, and 5.8% in education.

**Table 2.** Predicting LBC1936 phenotypes using methylation and genetic predictors for alcohol, BMI, smoking, and educational attainment

| Trait | DNAm Score | Polygenic Score | DNAm + Polygenic |
| --- | --- | --- | --- |
| BMI (kg/m<sup>2</sup>) | 12.2% | 10.3% | 19.5% |
| Alcohol (units per week) | 12.5% | 0.7% | 12.9% |
| Smoking (current/ever/never) | 60.6% | 2.8% | 61.0% |
| Educational attainment (years) | 2.6% | 4.0% | 5.8% |

For each trait, the proportion of phenotypic variance explained is presented for DNAm score, polygenic score and combined DNAm + polygenic scores.

**Figure 1.**
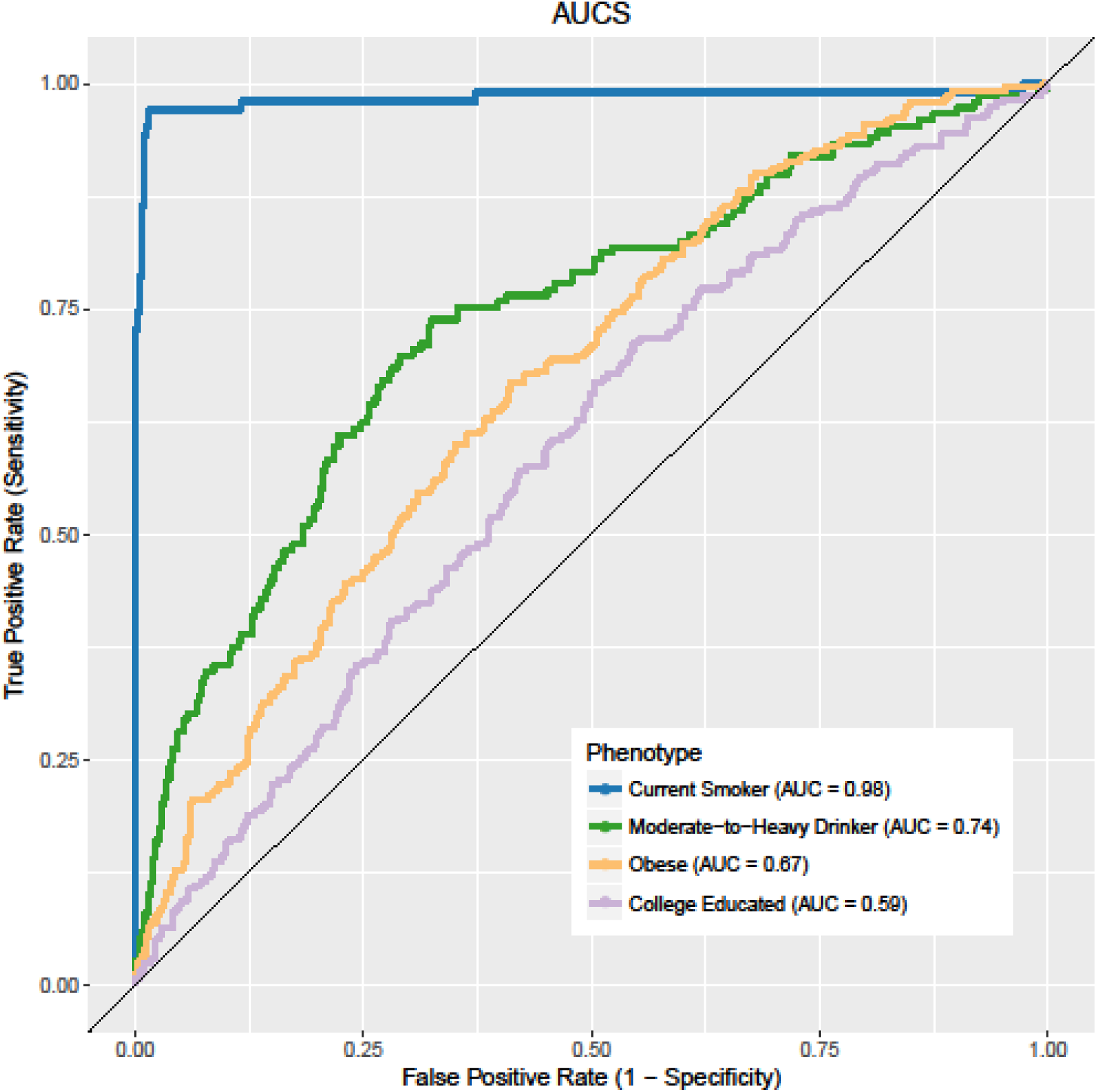
DNAm and polygenic prediction of alcohol, BMI, smoking, and educational attainment Proportion of phenotypic variance explained (R^2^) is plotted for four traits: BMI, smoking, alcohol and education based on each trait’s polygenic score (blue), DNA methylation-based score (green) and additive genetic + epigenetic score (orange).

### DNAm predictors classify phenotype extremes

For the Area Under the Curve (AUC) analyses that predicted the binary classified phenotypes in LBC1936, there were 652 controls and 242 cases for obesity, 755 light-to-moderate drinkers and 151 heavy drinkers, 423 non-smokers and 103 current smokers, and 233 and 671 individuals with >11 and ≤11 years of full-time education, respectively. There was near-perfect discriminatory power for the identification of current smokers (AUC=0.98), moderate discrimination of obesity from non-obesity (AUC=0.67) and of light-to-moderate drinkers from heavy drinkers (AUC=0.74), but only poor discrimination of those with more years of full-time education (AUC=0.59, **Figure 2**). Including the polygenic scores in addition to the DNAm predictors improved the prediction for obesity (AUC=0.71) and education (AUC=0.65) but not for the other traits. The smoking DNAm predictor was a significant addition to a logistic regression model for the binary education measure (smoking DNAm P=0.01, education DNAm P=0.08, and polygenic education P=2.5x10^−8^) but not for the models with obesity and moderate-to-high drinking as the outcomes (smoking DNAm P=0.31 and P=0.5, respectively).

**Figure 2.**
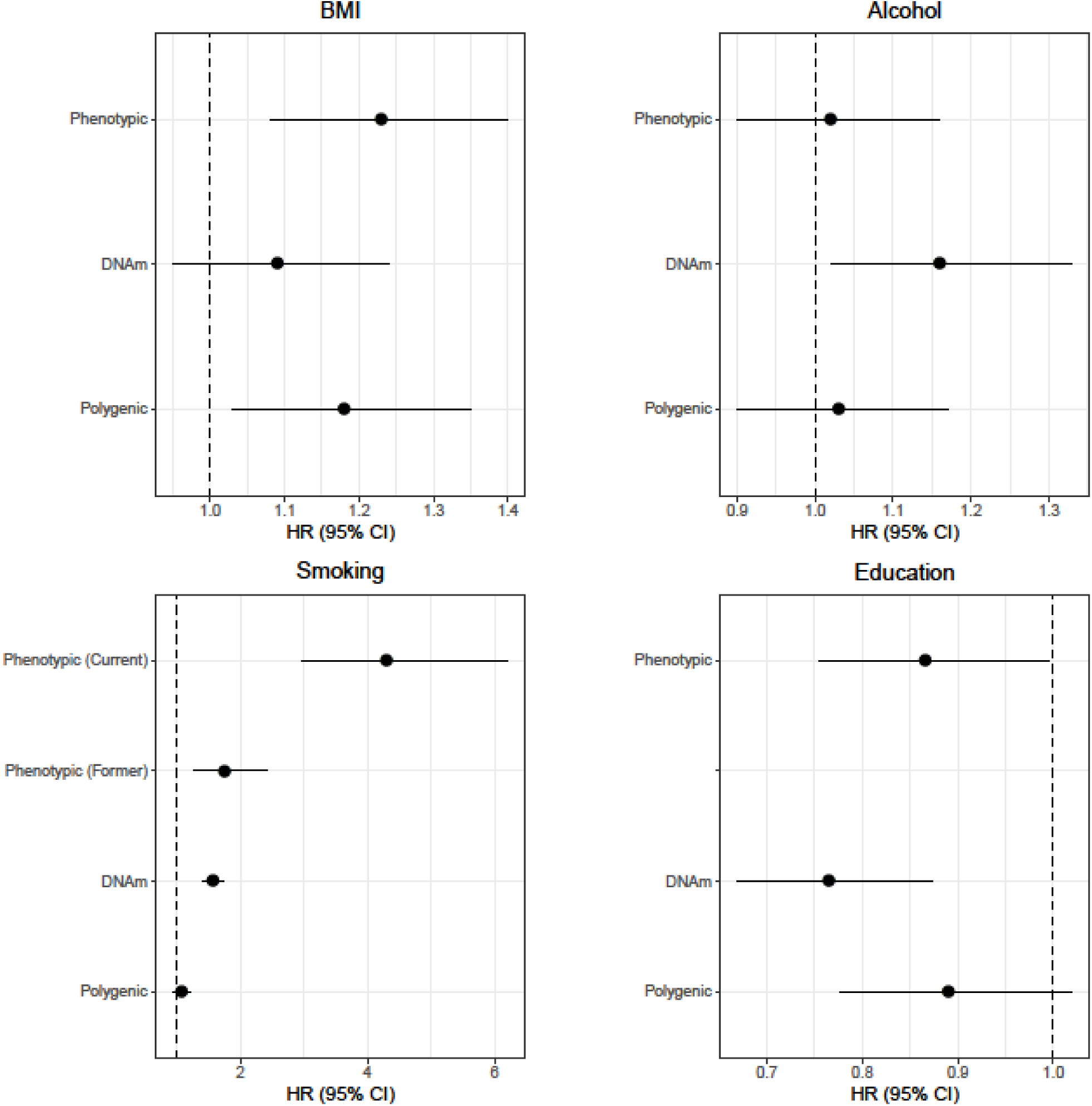
Receiver operating characteristic analysis for DNAm predictors of alcohol, smoking, and BMI. Shown are receiver operating characteristic curves for predicting moderate-to-heavy vs non-to-light drinkers, current smokers vs never smokers, obese vs non obese individuals, and high versus low-to-average education. Obese and non-obese are defined as BMI > 30 and ≤30kg/m^2^; moderate-to-heavy and non-to-light drinkers defined as drinking >21 and ≤21 units (men) or >14 and ≤14 units (women) of alcohol per week; highly educated individuals had >11 years of full-time education, compared to low-to-average education (≤11 years).

### DNAm predictors and mortality

Mortality in LBC1936 was assessed in relation to the phenotype, DNAm score, and polygenic score using Cox proportional-hazards models, adjusting for sex (**Table 3** and **Figure 3**). There were 214 deaths from 906 participants over 12 years of follow-up.

**Table 3.** Cox model outputs for phenotypic, DNAm and polygenic predictors of alcohol, smoking, education and BMI.

| **Trait** | **Predictor** | **HR** | **95% CI** | **P** |
| --- | --- | --- | --- | --- |
| Alcohol | Phenotypic | 1.02 | 0.90 – 1.16 | 0.73 |
|  | Epigenetic | 1.16 | 1.02 – 1.33 | **0.02** |
|  | Genetic | 1.03 | 0.90 – 1.17 | 0.68 |
| Smoking | Phenotypic (ex-smoker) | 1.76 | 1.28 – 2.42 | **4.5x10<sup>-4</sup>** |
|  | Phenotypic (current smoker) | 4.30 | 2.97 – 6.21 | **8.9x10<sup>-15</sup>** |
|  | Epigenetic | 1.57 | 1.40 – 1.75 | **1.0x10<sup>-15</sup>** |
|  | Genetic | 1.08 | 0.94 – 1.23 | 0.27 |
| Education | Phenotypic | 0.87 | 0.75 – 0.996 | **0.043** |
|  | Epigenetic | 0.76 | 0.67 – 0.87 | **7.8 x 10<sup>-5</sup>** |
|  | Genetic | 0.89 | 0.78 – 1.02 | 0.094 |
| BMI | Phenotypic | 1.23 | 1.08 – 1.40 | **0.001** |
|  | Epigenetic | 1.09 | 0.95 – 1.24 | 0.23 |
|  | Genetic | 1.18 | 1.03 – 1.35 | **0.016** |
| Alcohol | Phenotypic | 0.95 | 0.82 – 1.10 | 0.45 |
|  | Epigenetic | 1.19 | 1.03 – 1.38 | **0.02** |
| Smoking | Phenotypic (ex- smoker) | 1.44 | 1.01 – 2.05 | **0.042** |
|  | Phenotypic (current smoker) | 2.09 | 1.10 – 3.96 | **0.024** |
|  | Epigenetic | 1.31 | 1.08 – 1.60 | **0.006** |
| Education | Phenotypic | 0.90 | 0.79 – 1.04 | 0.15 |
|  | Epigenetic | 0.78 | 0.68 – 0.89 | **2.2x10<sup>-4</sup>** |
| BMI | Phenotypic | 1.21 | 1.06 – 1.39 | **0.006** |
|  | Epigenetic | 1.01 | 0.87 – 1.17 | 0.88 |

Cox proportional hazards outputs are presented for phenotypic, DNAm, and polygenic predictors of alcohol, smoking, education, and BMI in LBC1936. Each trait was assessed separately. Multivariate results are shown below the dashed line, assessing DNAm scores and their phenotypic analogues. Phenotypic never-smokers were used as a reference group for phenotypic ex-smokers and current smokers

**Figure 3.**
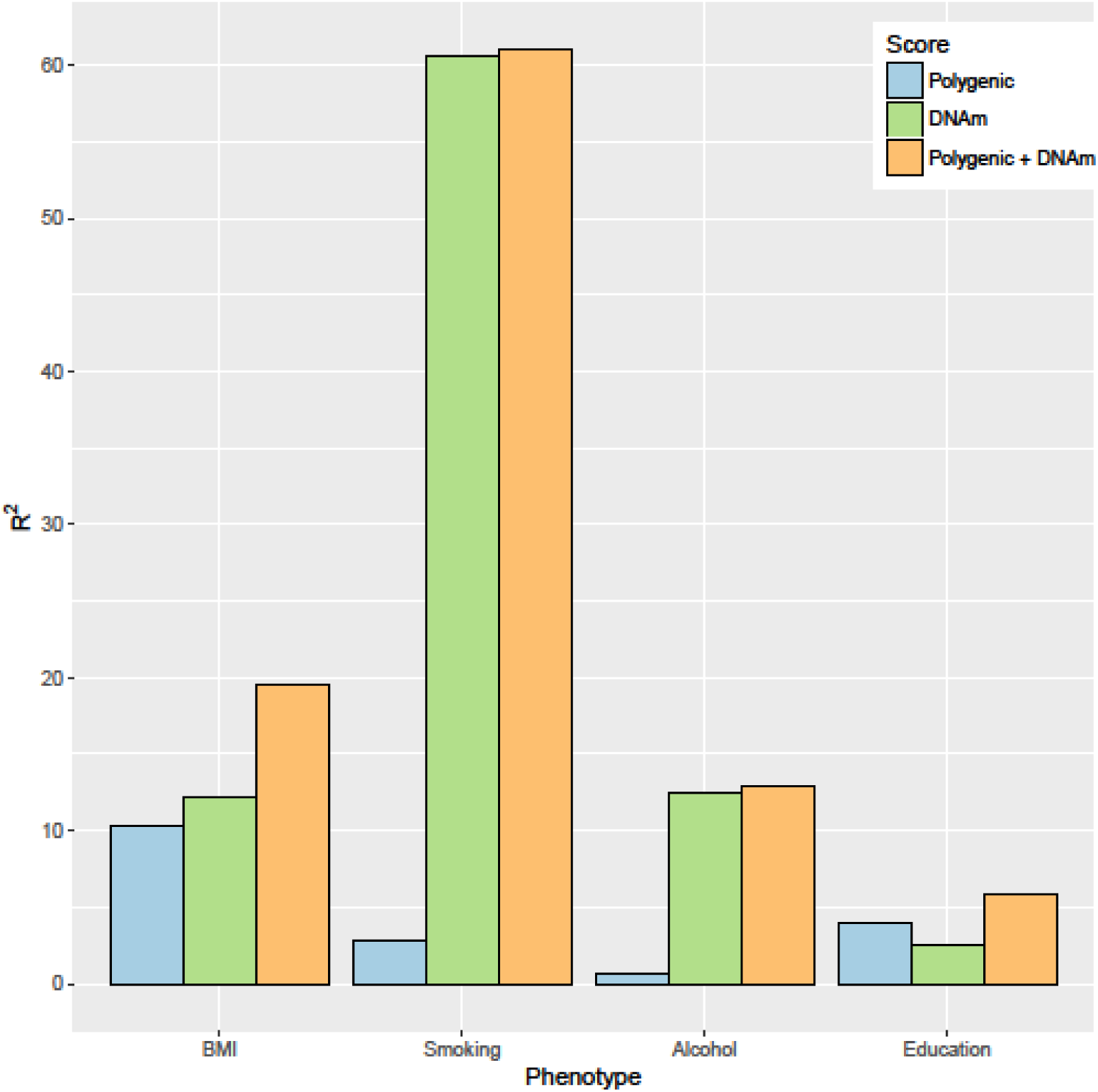
Hazard Ratios for phenotypic, epigenetic (DNAm), and genetic (polygenic) predictors of mortality. Forest plots show hazard ratios for phenotypic, DNAm and polygenic scores for BMI, alcohol consumption, smoking and education. Effect sizes are per standard deviation with the exception of phenotypic smoking, for which never smokers are used as a reference group. Horizontal lines represent 95% confidence intervals.

Higher phenotypic BMI and current/former smoking status (compared to never smokers) were associated with higher mortality risk (BMI: HR = 1.23 per SD, 95% CI = 1.08-1.40, P = 0.001; former smokers: HR = 1.76, 95% CI = 1.28-2.42, P = 4.5 x 10^−4^; current smokers: HR = 4.30, 95% CI = 2.97-6.21, P = 8.9 x 10^−15^). A mild protective effect was associated with higher educational attainment (HR = 0.87, 95% CI = 0.75-0.996, P = 0.043). No significant associations were observed in LBC1936 between risk of mortality and phenotypic alcohol consumption. A significant association was observed between mortality and the polygenic score for BMI (HR = 1.18, 95% CI = 1.03-1.35, P = 0.016) but not for the other three genetic scores. Higher mortality risk was associated with higher DNAm scores for smoking (HR = 1.57, 95% CI = 1.40-1.75, P = 1.0 x 10^−15^) and alcohol consumption (HR = 1.16, 95% CI = 1.02-1.33, P = 0.02), but not for BMI (HR = 1.09, 95% CI = 0.95-1.24, P = 0.23). A higher DNAm score for education was associated with lower mortality risk (HR = 0.76. 95% CI = 0.67-0.87, P = 7.8 x 10^−5^). The DNAm predictors of alcohol consumption and educational attainment were significantly associated with mortality following the addition of their phenotypic analogues to the respective models (DNAm HRs = 1.19 and 0.78, P = 0.02 and 2.2 x 10^−4^, respectively). The phenotypic and DNAm predictors of smoking were jointly associated with mortality (DNAm HRs = 1.31, 95% CI = 1.08-1.60, P = 0.006; phenotypic HR (former smokers) = 1.44, 95% CI = 1.01-2.05, P = 0.042; phenotypic HR (current smokers) = 2.09, 95% CI = 1.10-3.96, P = 0.024).

A final set of three survival models were considered. These covaried for the smoking DNAm predictor alongside the phenotype and DNAm predictor for (1) BMI, (2) alcohol, and (3) education (**Table 4**). Both the phenotypic BMI and smoking DNAm predictor were significant predictors of mortality (BMI HR = 1.28, 95% CI = 1.12-1.47, P = 3.0 x 10^−4^; DNAm smoking HR = 1.64, 95% CI = 1.46-1.84, P < 2.0 x 10^−16^). However, conditioning on the smoking DNAm predictor attenuated the association between the alcohol consumption and education DNAm predictors and mortality (P = 0.38 and P = 0.82, respectively). There were minimal differences to the effect sizes and interpretation of the findings after a further analysis that conditioned on white blood cell counts, with and without epigenetic scores for the corresponding trait and smoking DNAm score (**Supplementary Table 7**).

**Table 4.**
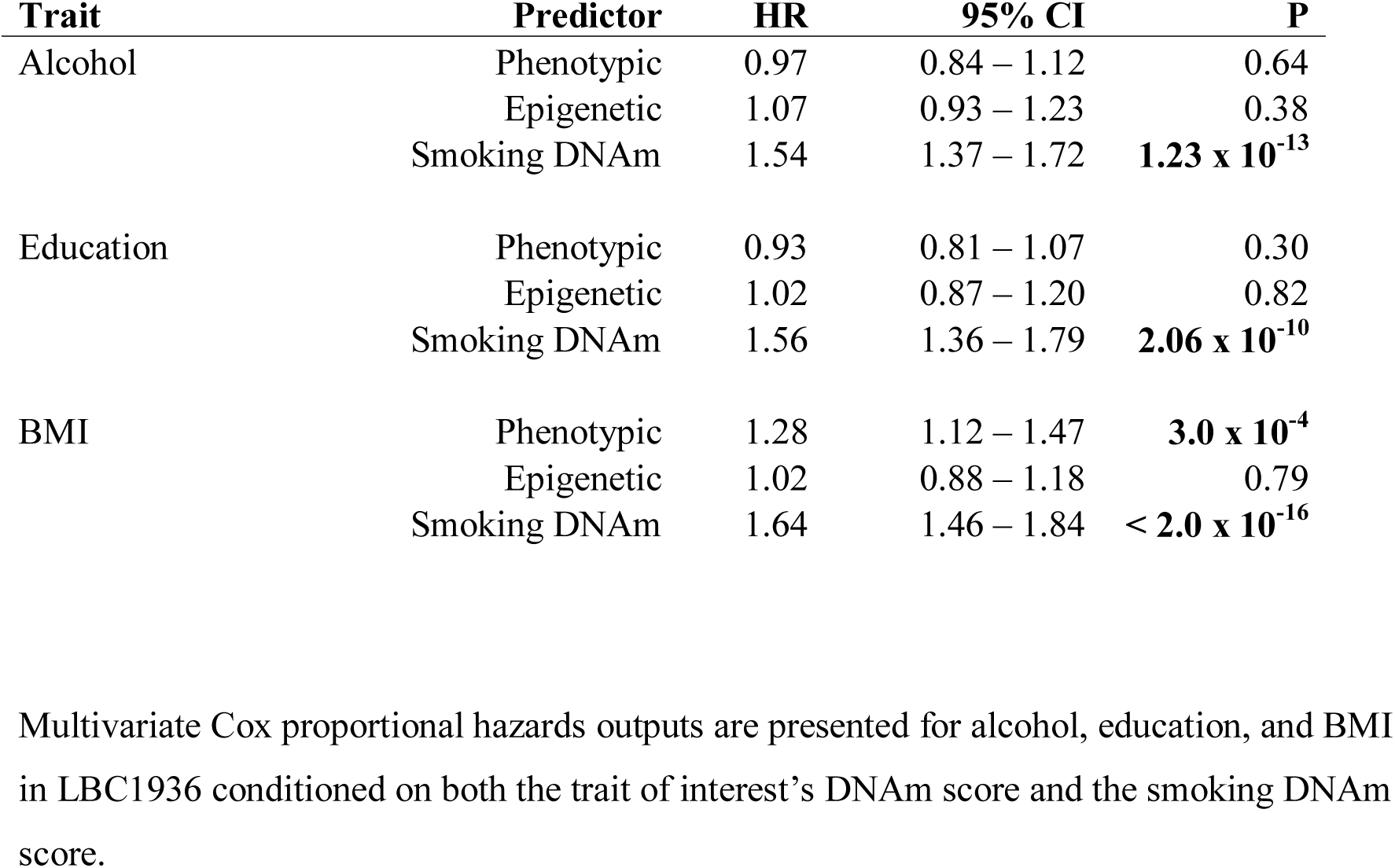
Multivariate Cox model outputs for alcohol, education, and BMI in LBC1936

| **Trait** | **Predictor** | **HR** | **95% CI** | **P** |
| --- | --- | --- | --- | --- |
| Alcohol | Phenotypic | 0.97 | 0.84 – 1.12 | 0.64 |
|  | Epigenetic | 1.07 | 0.93 – 1.23 | 0.38 |
|  | Smoking DNAm | 1.54 | 1.37 – 1.72 | **1.23 x 10<sup>-13</sup>** |
| Education | Phenotypic | 0.93 | 0.81 – 1.07 | 0.30 |
|  | Epigenetic | 1.02 | 0.87 – 1.20 | 0.82 |
|  | Smoking DNAm | 1.56 | 1.36 – 1.79 | **2.06 x 10<sup>-10</sup>** |
| BMI | Phenotypic | 1.28 | 1.12 – 1.47 | **3.0 x 10<sup>-4</sup>** |
|  | Epigenetic | 1.02 | 0.88 – 1.18 | 0.79 |
|  | Smoking DNAm | 1.64 | 1.46 – 1.84 | **< 2.0 x 10<sup>-16</sup>** |

Multivariate Cox proportional hazards outputs are presented for alcohol, education, and BMI in LBC1936 conditioned on both the trait of interest's DNAm score and the smoking DNAm score.

## Discussion

We have identified DNA methylation-based predictors for educational attainment, alcohol consumption, smoking, and BMI that: (1) explain varying degrees of proportions of their phenotypic variance, and do so independently from corresponding genetic predictors; (2) help to characterise individual differences; and (3) show clinical utility through prediction of mortality, and do so independently from phenotypic and genetic measures.

The DNAm predictors explained different proportions of the variance in the modifiable complex traits, from <3% for education to 12% for BMI and alcohol, and up to 60% for smoking. By combining genetic and epigenetic predictors we were able to augment these predictions to nearly 20% for BMI and 6% for education, whereas the alcohol and smoking predictions were largely driven by the DNAm predictors. The previous best estimate for genetic plus epigenetic BMI prediction was ~15% [10].

There is near-perfect discrimination between current and never smokers based on the smoking DNAm predictor and moderate discrimination between obese individuals and moderate-to-high drinkers. Differentiating those with a high level of education is more a function of genetics than DNAm, although the combined predictive power remains poor. Application of these predictors alongside existing DNAm-based age predictors [13, 14] may be of use in forensic investigations, given an unknown blood sample [15].

As with the previous EWAS analysis of education [5], there is a strong overlap with a smoking-related methylation signals. The strength of the correlation between the education and smoking DNAm predictors (r=-0.54) is particularly interesting when placed in context with their more modest phenotypic correlation (r=-0.14). Given that DNA methylation is highly predictive of smoking status [9], it may be the case that, should a single smoking-sensitive CpG feature in a DNAm predictor for another trait – here, education – then this drives a high correlation between the two DNAm predictors. It is of note that cg11902777, an established DNAm-based biomarker of smoking from the *AHRR* gene, was the feature with the fourth-largest coefficient in the education DNAm LASSO model. A DNAm education predictor excluding this feature/CpG correlated 0.98 with the primary predictor. Correlations between different CpG features within each of DNAm predictors may be responsible for the association observed between predictors.

The survival analysis in the out-of-sample prediction LBC1936 cohort yielded significant associations for the smoking, alcohol, and education DNAm predictors even after conditioning on their respective phenotypic values. When included as a covariate, the smoking DNAm predictor attenuated the DNAm – mortality associations for both the education and alcohol predictors. Consistent with our phenotype-based survival analyses, others have reported positive associations between mortality risk and both smoking and BMI [16, 17, 18] whereas higher educational attainment has been associated with a decreased mortality risk [19, 19]. A recent meta-analysis failed to find a significant relationship between alcohol consumption and all-cause mortality [21].

There are two key strengths to this study. First, the sample size of the Generation Scotland cohort, which is currently the single largest epigenetic epidemiology cohort in the world, enabled us to improve on previous DNAm predictors by: modelling all CpG sites simultaneously; training the predictor using cross-validation penalised regression modelling; and reducing heterogeneity in both phenotypic and methylation measurement through a single data collection and analysis protocol. Second, we could predict not only the relevant phenotypes of interest but also a clinically meaningful outcome (mortality) in our large, genetically homogenous, out-of-sample prediction cohort, LBC1936. Other studies with DNA methylation data and longitudinal disease follow-up for e.g., cardiometabolic, cardiovascular, and cancer-related outcomes will be able to further test the predictive performance of our DNAm predictors.

The Generation Scotland cohort contained related individuals who may be more phenotypically similar for the four traits under investigation. Residuals from sensitivity analyses that adjusted the phenotypes for pedigree structure as a random effect, in addition to age, sex, and population stratification as fixed effects, correlated highly (minimum Pearson r=0.96) to those from the models without pedigree adjustment. The older age range of LBC1936 and longitudinal follow-up enabled us to examine the ability of DNAm-based predictors for educational attainment, alcohol consumption, smoking and BMI to predict mortality, independently of the phenotypes themselves. As mentioned previously, the test cohort was older, had approximately 2 fewer years of education, were lighter drinkers, and heavier smokers relative to the training cohort. The DNAm predictors may perform differently on BMI, alcohol, smoking, and education measures in cohorts that are more analogous in age and phenotypic distribution to the training dataset, Generation Scotland.

## Conclusions

In summary, we showed that DNAm predictors are able to predict modifiable lifestyle factors with some success. They can also augment phenotypic prediction of mortality. Future studies should focus on other incident health outcomes, such as cardiometabolic disease and cancer. There is scope to use these DNAm predictors, in addition to DNAm-based predictors of age, to help identify lifestyle characteristics from DNA.

## Methods

### Training dataset for the DNAm predictors: Generation Scotland

The DNAm predictors were built on a subset of 5,100 individuals from Generation Scotland: the Scottish Family Health Study (GS:SFHS, hereafter abbreviated as GS), who had DNA methylation measured as part of a sub-study: Stratifying Resilience and Depression Longitudinally (STRADL). The parent cohort, GS, contains detailed cognitive, physical, health, and genetic data on over 22,000 individuals from across Scotland, aged between 18 and 99 years [22, 23]. It is a family structured, population-based longitudinal cohort study. Stored DNA samples from bloods collected at the study baseline (2006-2011) were used for the DNAm analysis.

### Methylation preparation in Generation Scotland

Quality control was performed on Illumina HumanMethylationEPIC BeadChip DNA methylation data from blood samples of 5,200 individuals from the Generation Scotland cohort. Briefly, visual inspection of a plot of log median intensity of methylated versus unmethylated signal [24] was used to identify outliers, which were excluded. Sample exclusions were also made where predicted sex, based on DNA methylation data, did not match the sex recorded in the GS database. Finally, samples were excluded if ≥1% of CpGs had a detection p-value in excess of 0.05. Probes were excluded if they had a beadcount below 3 in at least 6 samples and when ≥0.5% of samples had a detection p-value >0.05. This left data available for 860,926 methylation sites in 5,100 participants.

Further filtering was performed to remove (1) any sites with missing values, (2) non-autosomal sites, (3) non-CpG sites, and (4) CpG sites not present on the Illumina 450k array. Criterion (4) enables prediction into existing datasets as the majority of the CpG sites on the 450k array are present on the EPIC array.

### Phenotype preparation in Generation Scotland

We considered four phenotypes from Generation Scotland for the analysis: educational attainment, BMI, and self-reported alcohol consumption and smoking status. Educational attainment was measured via an ordinal scale: 0: 0 years, 1: 1-4 years, 2: 5-9 years, 3: 10-11 years, 4: 12-13 years, 5: 14-15 years, 6: 16-17 years, 7: 18-19 years, 8: 20-21 years, 9: 22-23 years, 10: ≥24 years of full-time education. It was treated as a continuous variable for the current analyses. BMI, assessed as the ratio of weight in kilograms to height in metres squared (kg/m^2^), was trimmed for extreme values (<17 and >50 kg/m^2^) before being log transformed. Alcohol was assessed in units per week and was only considered in those who reported that their intake as representative of a normal week. To reduce skewness in the distribution of alcohol consumption, a log(units + 1) transformation was performed. The addition of the constant retains non-drinkers, who reported a consumption of 0 units. Smoking was assessed in pack years (calculated by multiplying the number of packs smoked per day by the number of years the participant has smoked) for current and never smokers; ex-smokers were excluded due to complications in adjusting for time since cessation into the pack years calculation. Those who reported starting at age 10 or under were treated as data entry or self-report errors and were excluded along with non-smokers who had a non-zero (impossible) entry for pack years. As with the alcohol phenotype, a log(pack years + 1) transformation was used to reduce skew, leaving between 2,824 and 5,048 individuals, depending on phenotype (Table 1).

Each phenotype was then regressed on age, sex, and 10 genetic principal components [25] with the residuals being entered as the dependent variable in the LASSO models.

### LASSO regression in Generation Scotland

Penalised regression models were run using the glmnet library in R [26, 27]. 10-fold cross validation was applied and the mixing parameter (alpha) was set to 1 to apply a LASSO penalty. Coefficients for the model with the lambda value corresponding to the minimum mean cross-validated error were extracted and applied to the corresponding CpGs in an out of sample prediction cohort to create the DNAm predictors.

### The out-of-sample prediction cohort: Lothian Birth Cohort 1936

The Lothian Birth Cohort 1936 (LBC1936) [28, 29] was used for external DNAm predictions. LBC1936 is a cohort comprising individuals born in 1936, most of whom took part in the Scottish Mental Survey 1947. Participants were recruited to LBC1936 when they were aged approximately 70 years and have attended clinical examinations approximately every 3 years on up to 5 occasions. Detailed cognitive, physical, and health data have been collected, along with extensive ‘omics and biomarker data, including whole genome sequencing and longitudinal measures of DNAm, gene expression, and structural brain imaging. In the present study, DNAm was assessed in blood samples from wave 1 of the study between 2004 and 2007.

Mortality data in LBC1936 were obtained through data linkage to the National Health Service Central Register, provided by the General Register Office for Scotland (now National Records of Scotland). The mortality data used in the present analysis were correct as of January 2018.

### Methylation preparation in the Lothian Birth Cohort 1936

DNAm from whole blood was assessed in the Lothian Birth Cohort 1936 using the Illumina 450k methylation array. Over 90% of the 450k CpG sites are present on the EPIC array. Quality control details have been reported previously [30]. Briefly, after background correction, probes were removed if they were poorly detected (P>0.01) in >5% of samples or of low quality (via manual inspection). Samples were removed if they had a low call rate (P<0.01 for <95% of probes), a poor match between genotype and SNP control probes, or incorrect DNAm-predicted sex.

### Polygenic scoring in the Lothian Birth Cohort 1936

Polygenic scores were created in LBC1936 using PRSice [31] with clumping parameters of R^2^>0.25 over 250kb sliding windows. Genotyped data were generated at the Wellcome Trust Clinical Research Facility using the Illumina 610-Quadc1 array (San Diego). The SNP weights for all variants (P<1) for BMI [32], smoking [33], alcohol [34], and educational attainment [35] were taken from large genome-wide association studies (GWAS). Where LBC1936 was included in the discovery GWAS (educational attainment [35]), the meta-analysis was re-run after its exclusion.

### Phenotypes in the Lothian Birth Cohort 1936

DNAm predictors for smoking, alcohol, BMI, and educational attainment were used to explain variance in their corresponding phenotypes in the out-of-sample cohort, LBC1936. Phenotype measurement details in LBC1936 are as follows: self-reported smoking status (current smoker, ex-smoker, never smoked), alcohol consumption in a typical week (recoded into units), and education (years of full-time education) were assessed along with BMI (defined as the ratio of weight in kilograms divided by height in metres squared). Binary categorisations of smoking (current versus never), BMI (>30kg/m^2^ versus ≤30kg/m^2^, defined as obese and non-obese, respectively), education (>11 years versus ≤11 years, which is roughly equivalent to a college education level for LBC1936), and alcohol consumption were used as outcomes for receiver operating characteristic curve analyses. Sex-specific dichotomisations were applied to the alcohol consumption phenotype, as per UK health recommendations at the time of data collection (≤21 units per week versus >21 units per week for males, and ≤14 units per week versus >14 units per week for females; corresponding to moderate and heavy alcohol consumption in each gender, respectively.

### Prediction Analysis in the Lothian Birth Cohort 1936

There were four main aims for the prediction analysis: (1) to identify the proportion of phenotypic variance explained by the corresponding DNAm predictor; (2) to determine if this was independent of the polygenic (genetic) signal for each phenotype; (3) to obtain area under the curve (AUC) estimates for binary categorisations of BMI, smoking, alcohol consumption, and college education; and (4) to identify if the phenotype, polygenic score, or DNAm predictor explained mortality risk and if they do so independently of one another.

Linear regression models were used to explore aims (1) and (2). Ordinal logistic regression was used for the categorical smoking variable (never, ex, current smoker). Age and sex were considered as covariates, the phenotypic measure was the dependent variable, and the polygenic score or DNAm predictor were the independent variables of interest. Incremental R^2^ estimates were calculated between the null model and the models with the predictors of interest. An additive genetic and epigenetic model for BMI in the Lothian Birth cohort 1936 has been reported previously, although a different DNAm predictor, based on unrelated individuals, was derived from the Generation Scotland data [36]. Receiver operating characteristic curves were developed for smoking status, obesity, high/low alcohol consumption, and college education and areas under the curve were obtained using the pROC library in R [37](Aim 3). Finally, Cox proportional hazards survival models [38] were used to examine the associations with mortality listed under Aim 4. Sex was included as a covariate in all models.

## Declarations

### Authors’ Contributions

Conception and design: REM

Data analysis: DLM, REM

Drafting the article: DLM, REM

Data preparation: DLM, SJR, RMW, SWM, AFM, and REM

Data collection: ADM, DJP, NRW, PMV, AMM, KLE, IJD

Revision of the article: all authors

### Availability of Data and Material

Generation Scotland and Lothian Birth Cohort data are available upon request. Please contact

AC or IJD or visit the study websites:

http://www.generationscotland.co.uk/

https://www.lothianbirthcohort.ed.ac.uk/content/collaboration

### Ethics Approval and Consent to Participate

All components of GS:SFHS received ethical approval from the NHS Tayside Committee on Medical Research Ethics (REC Reference Number: 05/S1401/89). GS:SFHS has also been granted Research Tissue Bank status by the Tayside Committee on Medical Research Ethics (REC Reference Number: 10/S1402/20), providing generic ethical approval for a wide range of uses within medical research.

Ethical permission for the LBC1936 was obtained from the Multi-Centre Research Ethics Committee for Scotland (MREC/01/0/56) and the Lothian Research Ethics Committee (LREC/2003/2/29). Written informed consent was obtained from all participants.

### Consent for Publication

Not applicable

## Acknowledgements

Not applicable

